# On the stabilization of plant lipid droplets: Dynamic interplay between oleosins and phospholipids

**DOI:** 10.64898/2025.12.09.693226

**Authors:** Xuefeng Shen, Ivan Alcazar Salazar, Xingfa Ma, Ketan A. Ganar, Zohaib Hussain, Emmanouil Chatzigiannakis, Constantinos V. Nikiforidis, Jasper van der Gucht, Siddharth Deshpande

## Abstract

As the central organelles of lipid and energy homeostasis in plant seeds, lipid droplets (LDs) consist of a neutral lipid core, decorated by phospholipids (prominently phosphatidylcholines, PCs) and surfactant-like proteins (mostly oleosins, OLs). So far, the dynamic interplay between PCs and OLs at the LD interface remains unclear. The presented work addresses this knowledge gap by reconstituting oil-in-water emulsions stabilized by OLs and PCs using microfluidic systems. Our results show that the resistance to droplet coalescence is primarily provided by OLs. We further reveal that the addition of PCs alters the assembly of OLs at the interface, reducing the OL network density and interfacial elasticity, thereby rendering a weaker interface. In conclusion, our study suggests complementary roles, with OLs acting as the primary stabilizers while PCs playing a destabilizing role. This contrast likely contributes to the observed metastability of LDs and can be exploited to design stimuli-responsive emulsions.

**HIGHLIGHTS:**

- Oleosins form an interfacial network that stabilizes the oil-water interface.
- Phosphatidylcholines globally weaken the oleosin network and promote droplet coalescence.
- Microfluidics enables controlled reconstitution and real-time analysis of lipid droplets.
- Oleosin–phospholipid interplay explains lipid droplet metastability and can guide the design of bio-inspired emulsions.

## INTRODUCTION

Emulsions are metastable colloidal systems composed of two immiscible fluids, with one fluid phase dispersed within the other, stabilized by interfacial agents such as surfactants, amphiphilic polymers, or particles. In addition to their widespread application in various industries such as pharmaceuticals, paints, foods, healthcare, and oil recovery^1^, emulsions are also ubiquitous in biological systems, such as milk, where fat globules are dispersed in the aqueous phase^2^, lipid droplets (LDs) stabilized by bile salts in digestive system^3^, and LDs within plant and mammalian cells^4^. LDs in plants, also known as oleosomes, represent the dispersed phase of oil-in-water emulsions within the aqueous cytosol of plant cells. LDs exhibit a unique architecture, comprising a hydrophobic core of neutral lipids, most commonly triacylglycerols (TAGs) and sterol esters, surrounded by a dense phospholipid monolayer^5^. Besides the phospholipids, the LD membrane is decorated with various proteins, among which oleosins (OLs) are the predominant type^6^. Oleosins exhibit two hydrophilic N- and C-terminal that extend outside or on the phospholipid monolayer, and a central hydrophobic domain with a proline knot that penetrates into the TAG core like an anchor^6^, as schematically shown in Figure 1A. Oleosins are crucial for the stability of LDs, particularly during seed desiccation, rehydration, and environmental stresses such as temperature fluctuations^7^. The phospholipids that compose the LD monolayer also function as natural surfactants, preferentially adsorbing to the interface due to their amphiphilic nature. Both these molecules reduce interfacial tension, confer bending elasticity to the monolayer, and thereby enhance the stability of LDs against coalescence^6^.

**Figure 1:**
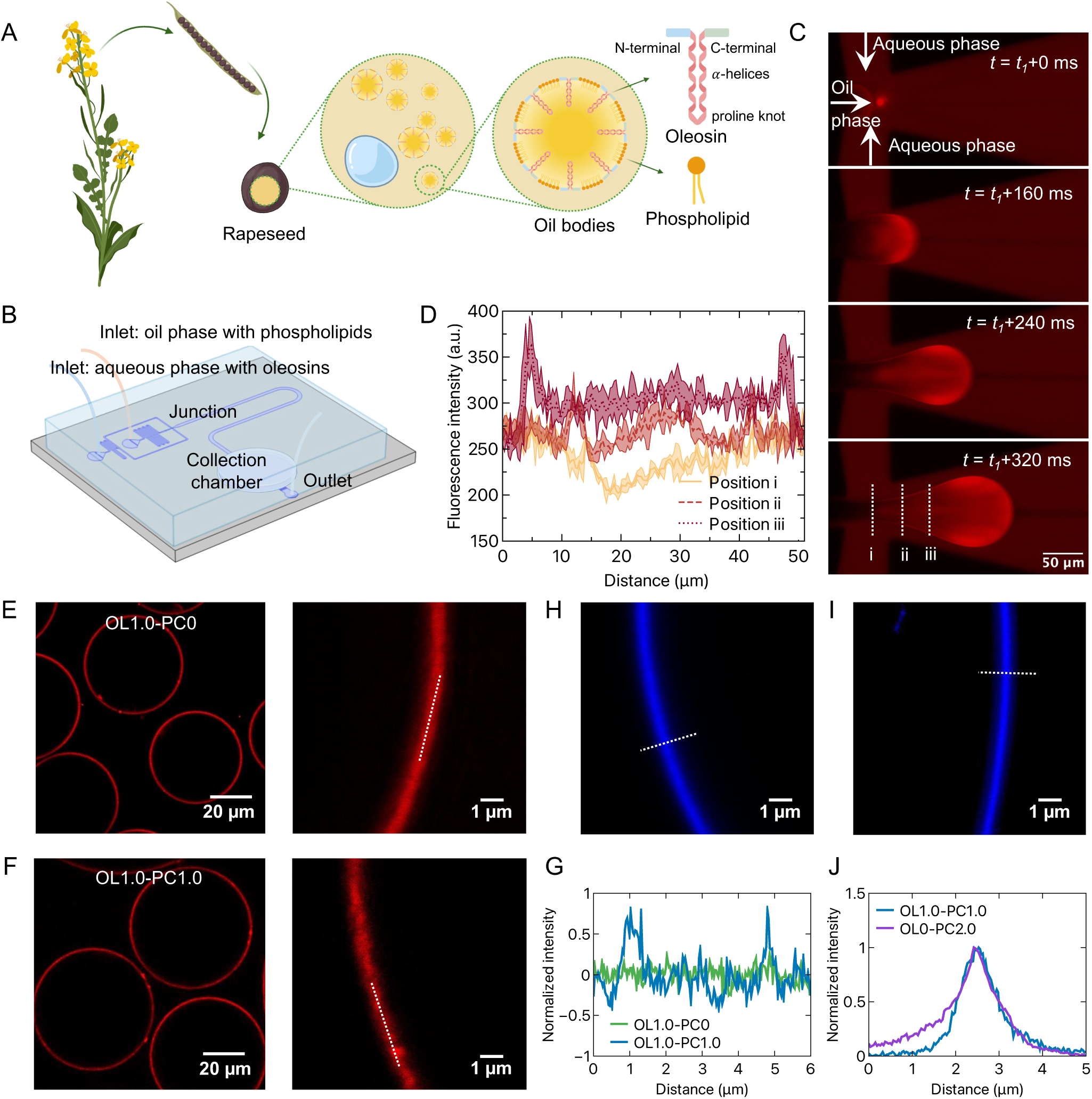
Reconstitution of LDs using microfluidics. **A** Schematic representation of an oilseed plant, the source of OLs. OLs are localized within the seeds and, together with phospholipids, act as interfacial stabilizers for TAG droplets dispersed in the aqueous cytosol. **B** Schematic of the microfluidic device, illustrating the inlets for the OL dispersion and the oil phase, the flow-focusing junction, a downstream protein adsorption channel, a collection chamber, and an outlet. **C** Fluorescence microscopy images showing rapid OL adsorption at the droplet interface prior to detachment from the junction. **D** Corresponding fluorescence intensity profiles across different regions of the droplet. **E** STED micrographs of collected droplets showing OL localization at the droplet interface without PCs present (left), with a magnified view (right). **F** STED micrographs of collected droplets showing OL localization at the droplet interface with PCs present (left), with a magnified views (right). **G** Normalized intensity plots corresponding to interfaces stabilized solely by OLs and stabilized by both OLs and PCs show that the interface containing only OLs appears smoother. **H-I** Confocal micrographs showing the localization of fluorescently labeled phospho-lipids at the droplet interface in the presence (**H**) and absence (**I**) of OLs. **J** The corresponding normalized intensity profiles across the interfaces indicate that interface containing only phospholipids is thinner. The OL concentration is 1.0 mg/mL in panels (**C–H**), and PC is additionally present at 1.0 mg/mL in panels **F** & **H** and at 2.0 mg/mL in panel **I**.

In nature, plant seeds undergoing dormancy or germination are exposed to a wide range of physicochemical conditions in the soil^8^. LDs within these seeds—particularly in salt- and alkali-tolerant species—exhibit remarkable resistance to fluctuations in soil salinity and pH, thereby contributing to seed viability under harsh environmental conditions^9,10^. These LDs remain discrete and resist aggregation or coalescence, even under extreme environmental stresses such as moisture and temperature fluctuations during seed maturation. Notably, they also retain their structural integrity and stability after isolation from seed cells and subsequent suspension in aqueous solution^11^. The remarkable stability of LDs has drawn increasing attention to their interfacial OLs and phospholipids in recent years^12–14^, as there is a growing interest in replacing synthetic emulsifiers with natural alternatives and substituting animal-based ingredients with functional plant-based counterparts^15,16^. However, with the complex cytoplasmic environment surrounding the LDs and with both phospholipids and OLs displaying interfacial activity, the precise interfacial structure, the mechanism of interfacial adsorption, and the interaction between PCs and OLs at the interface and their respective roles in stabilization remain unclear. Previous work has focused on the effects of phospholipids and OLs on the interfacial rheological properties^14^; however, it disregards the stability of the LDs under changing environmental conditions or external forces, while the insight into the mechanisms underlying the droplet physical stability are still missing. To obtain a better understanding, we investigated a minimal reconstituted system comprising key components of LDs and allowing analysis of the interface in a controllable way.

While conventional emulsification methods produce polydisperse droplets that impede systematic analysis^2^, microfluidic emulsification enables precise control over droplet size and interfacial composition, and also facilitates *in situ* microscopic observation^17,18^. Beyond the diffraction limitations of standard light microscopy and the invasive sample preparation required for electron microscopy, super-resolution techniques overcome these constraints, providing nanometric, multicolor imaging of interfacial protein structures^19^. In addition to microscopic observation, the thin film balance (TFB) is a powerful tool for investigating droplet physical stability, enabling controlled studies of the stabilization mechanisms associated with various interfacial stabilizers^20–23^. In this study, we utilized the above-mentioned techniques to gain insight into the respective roles of OLs and PCs in stabilizing the LD interface. We used on-chip microfluidics to generate model LDs stabilized via OLs and PCs, at varying ratios. Our on-chip setup enabled convenient droplet collection, exchange of the continuous phase, and real-time visualization of droplet coalescence behavior. We demonstrate that the coalescence resistance of the oil droplets in response to variations in pH and ionic strength is primarily attributed to the presence of OLs. PCs, in contrast, do not contribute to droplet stability across a wide pH range and even promote coalescence under increased ionic strength. Stimulated emission depletion (STED) microscopy results show that the adsorbed OLs assemble into a network-like structure at the interface, and a significantly reduced network density is observed upon the addition of PCs. Furthermore, interfacial rheology reveals that the addition of PC reduces the elastic modulus of an OL-stabilized interface. The drainage and rupture process of thin films indicate that interfaces stabilized by both OLs and PCs exhibit lower resistance to an applied pressure step, compared to those stabilized solely by OLs. Collectively, these findings reveal that OLs are superior interfacial stabilizers compared to PCs, at least in simplified oil droplet systems. Conversely, it also brings up the intriguing possibility that PCs might actively destabilize the LDs when required, thus making the two stabilizers work in tandem.

## RESULTS

### 0.1 *In vitro* reconstitution of LDs using on-chip microfluidics

OLs dispersed in the aqueous phase were extracted from rapeseeds following our recently published protocols^24^ (see Methods for details). SDS-PAGE analysis of the protein composition showed a predominant band at approximately 17 kDa as expected (Supplementary Figure 1). Phosphate-buffered saline (PBS) was used to disperse the OLs. The pH of the solution was adjusted to 2 using 1 M HCl, as the resulting high negative surface charge provides sufficient electrostatic repulsion between OLs, thereby improving their dispersibility^13^. The size distribution of OLs dispersed in an aqueous system (pH 2) was characterized using dynamic light scattering, which gave a broad size distribution ranging from 20 to 200 nm, with a primary peak around 50 nm (Supplementary Figure 2), indicating that they are present as micelles or micellar aggregates^13^. PCs, which account for approximately 60% of phospholipids in natural LDs^25^, were selected and dispersed in sunflower oil (see Methods for details). Next, we developed a lab-on-a-chip microfluidic platform to facilitate the efficient reconstitution, collection, and imaging of LDs. A schematic of the device is presented in Figure 1B. The chip was fabricated using a standard soft lithography technique to create a multi-height polydimethylsiloxane (PDMS)-based microfluidic device (see Methods for fabrication details). In a typical experiment, sunflower oil droplets (with or without PCs) were generated at the flow-focusing junction and transported through a down-stream adsorption channel, where they encountered the oleosin-containing aqueous phase. To ensure that the droplets were initially less compressed physically during the analysis, the collection chamber height was kept at 40 *µ*m for all devices (Supplementary Figure 3).

Although the adsorption of OLs at the oil–water interface is generally thermodynamically favorable due to their amphiphilic nature, their role as interfacial stabilizers in microfluidic systems remains insufficiently understood. Emulsions are metastable dispersions, and the stability of oil droplets is strongly influenced by the OL adsorption at the oil–water interface. To verify OL adsorption, we fluorescently labeled OLs (see Methods for details) and mixed them with unlabeled OLs at a 1:10 molar ratio to generate droplets within the microfluidic device. As shown in Figure 1C, oleosins rapidly adsorbed at the interface prior to the droplet pinch-off at the junction. This spontaneous and rapid interfacial adsorption was further confirmed by fluorescence intensity profiles across different regions of the forming droplet (Figure 1D). The early adsorption of OLs provides initial protection against coalescence, thereby enabling the collection of stable droplets downstream.

Next, we took advantage of the on-chip settings to replace the continuous phase containing fluorescently labeled OLs with a clean, protein-free solution, thus minimizing background fluorescence. In these conditions, we employed super-resolution (STED) microscopy to take a closer look at the oil-water interface stabilized solely by OLs (Figure 1E) as well as by both OLs and PCs (Figure 1F), a comparison between confocal and STED micrographs is shown in Supplementary Figure 4. When stabilized solely by OLs, fluorescent proteins were observed exclusively at the droplet interface, with no signal detected in the droplet interior or in the surrounding medium, confirming their strong interfacial localization. When the LDs were stabilized by both OLs and PCs, strong OL fluorescence was still obtained at the interface, indicating that despite competitive adsorption by PCs, a significant amount of OLs still adsorbed at the interface. However, as can be seen from the magnified images, OL fluorescence at the interface appeared rougher when compared to the interface when stabilized solely by OLs, suggesting that PCs altered the spatial arrangement of OLs at the droplet interface. Figure 1G compares the normalized intensity profiles of two interfaces—one adsorbed with only OLs and the other with both OLs and PCs, respectively, making it clear that the interface stabilized solely by OLs appears smoother. Use of fluorescently labeled phospholipids (18:1 Liss Rhod PE, PC:Rhod-PE = 10:1, molar ratio) together with confocal microscopy confirmed the presence of PCs at the interface, shown in Figure 1H, further confirming that the reconstituted LDs were co-stabilized by both OLs and PCs. Furthermore, the confocal micrograph (Figure 1I) and the corresponding fluorescence intensity profile (Figure 1J) show that the PC-only interface appears thinner than the OL-containing interface. This observation suggests that the rougher interface containing both OLs and phospholipids may adopt a multilayer-like structure in which phospholipids can reside.

To assess the effect of OLs and PCs on reconstituted LD monodispersity, we compared the size distributions of droplets generated at varying PC concentrations while maintaining a constant OL concentration (Figure 2A). The histograms show four batches of droplets with radii ranging from 40 to 50 *µ*m. The coefficient of variation (CV) ranged from 4 to 8% of the mean for the first three cases, indicating a relatively monodisperse size distribution. In contrast, CV value increased to 15% when PC concentration was highest. Micrographs of collected LDs without PCs and with the highest PC concentration are shown in Figure 2B and C, respectively. Figure 2C clearly shows unusually large droplets, likely resulting from coalescence. This coalescence primarily occurred upstream in the adsorption channel, while droplets were still flowing through the adsorption channel. When interfacial stabilizers fail to provide adequate protection, droplets may merge upon collision with neighboring droplets, as illustrated in Figure 2D, leading to the formation of larger droplets. These findings suggest that, contrary to common assumptions^26^, phospholipids may not necessarily improve droplet stability in combination with OLs. The observed coalescence of droplets stabilized by OLs and PCs is noteworthy, given that phospholipids are frequently employed as interfacial stabilizers in emulsions^26^. Based on these results, we set out to study the effects of OLs and PCs on coalescence resistance under varying pH, ionic strength, and mechanical force, as well as their influence on interfacial properties.

**Figure 2:**
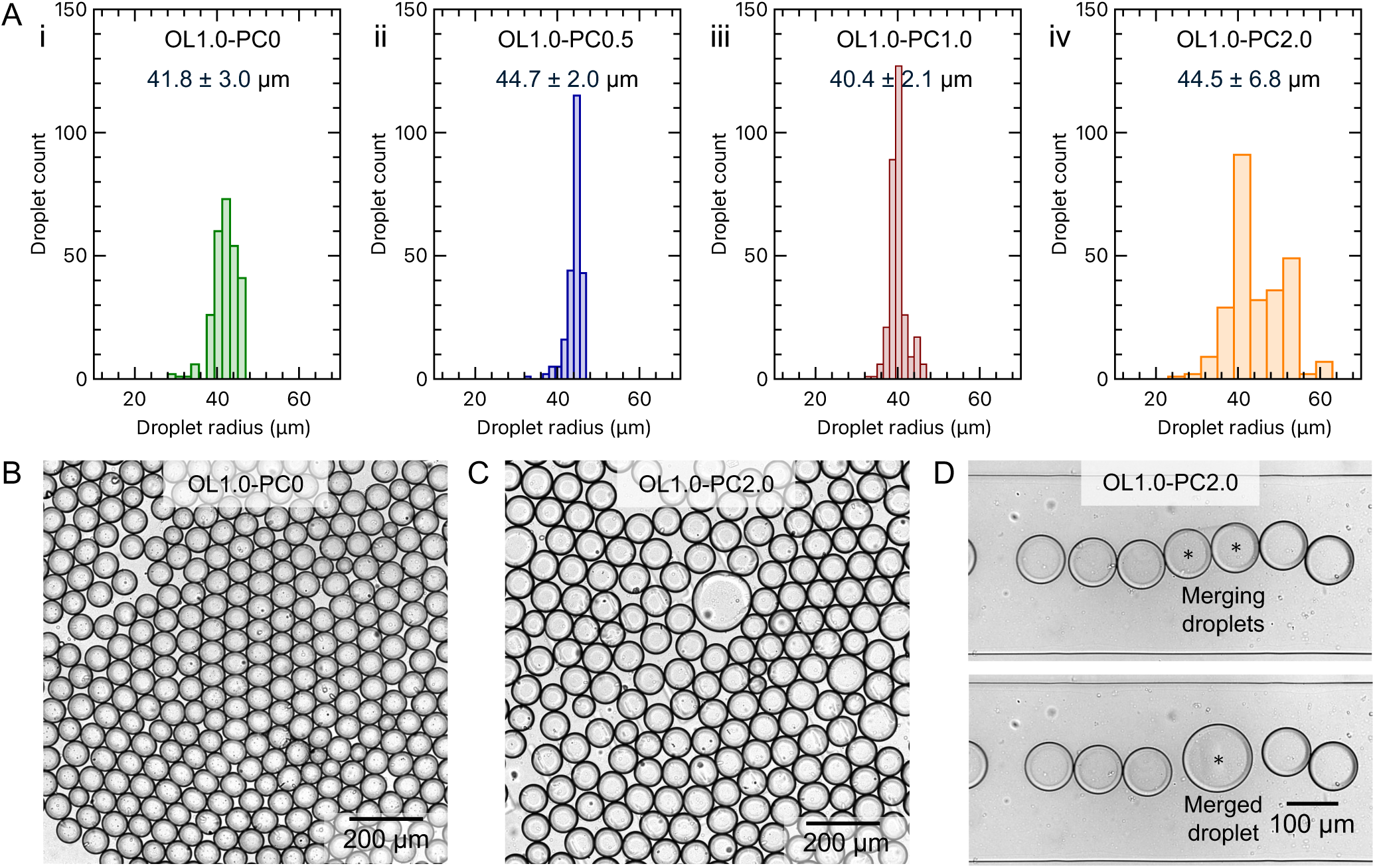
Effect of OL/PC mass ratio on the monodispersity of reconstituted LDs. **A** Histogram showing the size distribution and monodispersity of reconstituted LDs stabilized by OLs and PCs at varying concentrations. The OL concentration is fixed at 1.0mg/mL in all cases, while (i), (ii), (iii), and (iv) correspond to PC concentrations of 0 mg/mL, 0.5 mg/mL, 1.0 mg/mL, and 2.0 mg/mL, respectively. **B** Micrograph of collected LDs stabilized by OLs. **C** Micrograph of collected LDs stabilized by both OLs and PCs. Droplets appear slightly larger and less monodisperse compared to those stabilized solely by OLs. **D** Upon the addition of PC to the oil phase, droplets more frequently exhibit coalescence behavior in the adsorption channel; coalesced droplets are marked with a star.

### 0.2 Effect of pH and salt stress on OL/PC-coated LDs

Inspired by biological observations that LDs in plant seeds exhibit remarkable stability across a wide range of physicochemical conditions, we sought to determine which of the two primary interfacial stabilizers—OLs or PCs—plays a more dominant role in stabilizing LDs. Understanding their relative contributions is crucial for optimizing their use in the formulation of plant-protein-based emulsions, which often encounter environmental stresses during industrial processing, such as elevated temperatures, pH fluctuations, and changes in ionic strength. Having established a microfluidic approach for the reconstitution and collection of LDs, we further investigate their stability in response to variations in the pH and ionic strength of the continuous phase. Figure 3A illustrates a typical coalescence experiment involving a batch of oil droplets in the collection chamber. A new continuous phase at pH 5 is introduced from the top, replacing the original continuous phase at pH 2 used during droplet production. It took approximately 10 s for the continuous phase within the observed region (1100 × 1100 *µ*m) to be fully replaced. Initially, the droplets remained stable; however, upon arrival of the new continuous phase—as indicated by the orange arrows—some droplets began to coalesce, while others remained stable until the new phase reached their location. A 10-min video was recorded continuously until no further coalescence events were observed. We quantified the percentage of coalescence events over 10 minutes (see Methods for details).

**Figure 3:**
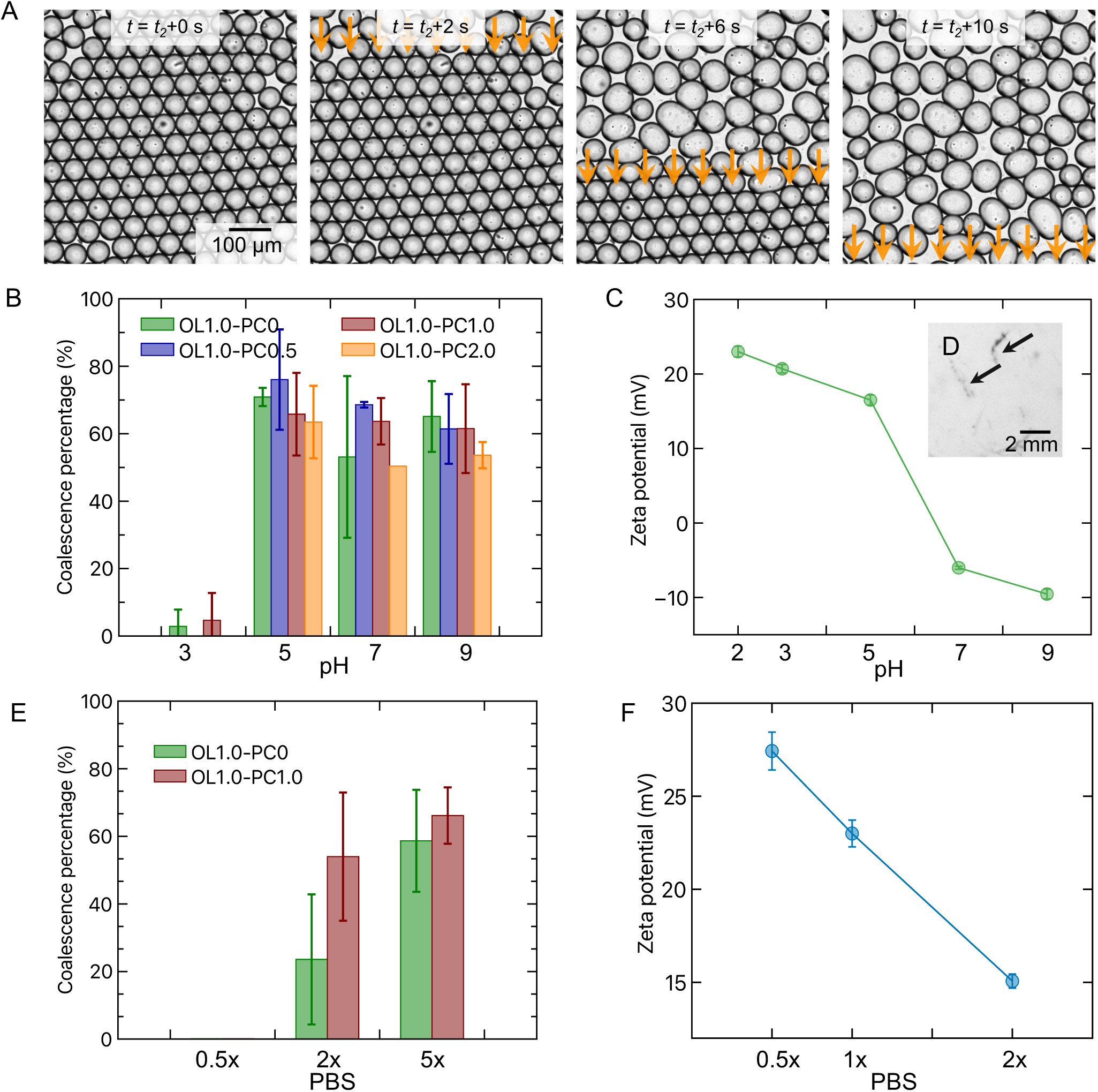
The role of OLs and PCs on the coalescence stability of LDs under varying pH and ionic strength. **A** Time-lapse microscopy images illustrating the coalescence assay. Tightly packed LDs within the collection chamber coalesce to varying degrees upon the introduction of a new continuous phase (pH 5 in this case), which replaces the initial pH 2 solution. Orange arrows indicate the direction from which the continuous phase is replaced. The droplets are stabilized solely by OL at a concentration of 1.0 mg/mL. **B** Coalescence percentage of OL-stabilized LDs with varying PC concentrations at different pH values. All formulations exhibited high stability at pH 3, but the coalescence percentage increased drastically at higher pH values. Although not statistically significant, droplets with the highest PC concentration (2.0 mg/mL) showed a trend toward lower instability across pH 5, 7, and 9. Data are presented as mean ± standard deviation; *N* = 3. **C** Zeta potential of OLs in water as a function of pH, showing a progressive decrease in surface potential with increasing pH. **D** Visual evidence of OL precipitation at pH 5, which is near its isoelectric point, in the bulk phase. **E** Coalescence percentage of LDs stabilized by OL alone (OL1.0–PC0) or with added PC (OL1.0–PC1.0) under different ionic strengths (0.5×, 2×, and 5× PBS). At low ionic strength (0.5× PBS), droplets remained stable; however, increasing the ionic strength led to greater coalescence, with the addition of PC offering no improvement—and potentially reducing—stability under high-salt conditions. Data are presented as mean ± standard deviation; *N* = 3. **F** Zeta potential of OLs in water as a function of PBS concentration, illustrating a reduction in surface potential with increasing ionic strength.

We first evaluated the effect of pH on the coalescence stability of LDs stabilized by OLs with varying PC concentrations. This investigation aimed not only to understand how OLs and PCs in seeds protect LDs from coalescence under varying pH conditions (pH values of 3, 5, 7, and 9) but also to assess their effectiveness as natural emulsifiers in resisting pH-induced destabilization of emulsions and foams. As shown in Figure 3B, all formulations exhibited minimal coalescence at pH 3, indicating strong droplet stability under acidic conditions, due to strong electrostatic repulsion. However, coalescence increased markedly at pH 5. Zeta potential measurements of OLs in water at different pH values with a constant PBS concentration, shown in Figure 3C, reveal a gradual decrease in surface potential with increasing pH. This suggests that the isoelectric point (pI) of OLs used in this study is approximately at pH 6. Near the pI, protein solubility decreases sharply, leading to visible precipitation (Figure 3D). This behavior likely arises from the loss of electrostatic repulsion between OLs due to a reduction in the net surface charge, primarily provided by the charged amino acid residues within the N- and C-termini. As electrostatic repulsion diminishes, attractive interactions—such as hydrophobic and van der Waals forces—become dominant, promoting protein aggregation. Although this phenomenon is observed in the bulk phase, a similar interfacial transition is expected, weakening interfacial strength and enhancing droplet coalescence. A decrease in electrostatic interactions between neighboring droplets with protein-rich interfacial layers is also anticipated. Coalescence occurs when the applied or internal liquid pressure (*P_L_*) exceeds the sum of the capillary pressure (Π*_st_*) and the electrostatic repulsive pressure (Π*_el_*), offset by the attractive van der Waals pressure (Π*_vdW_* ), according to the criterion *P_L_ >* Π*_st_* + Π*_el_ −* Π*_vdW_ .* Across pH 5, 7, and 9, PC concentration showed no significant effect on the coalescence stability of OL-stabilized droplets, indicating that the observed variations between formulations were not statistically significant and that PC provided no substantial improvement in pH resistance under these conditions.

Figure 3E presents the coalescence percentages of LDs stabilized by 1.0 mg/mL OL, with or without 1.0 mg/mL PC, under varying ionic strengths. The ionic strength was adjusted to 0.5×, 2×, and 5× the standard PBS concentration . At low ionic strength (0.5× PBS), no measurable coalescence was observed, indicating that electrostatic repulsion was sufficient to maintain droplet stability. However, as ionic strength increased, coalescence also increased for both formulations, leading to reduced electrostatic stabilization due to charge screening. Notably, at 2× and 5× PBS, the OL1.0–PC1.0 formulation exhibited greater coalescence than OL1.0–PC0, suggesting that the addition of PC did not enhance—and even impairs—interfacial stability at high-salt conditions. This is somewhat surprising, as one would expect the addition of zwitterionic PC to improve stability against salt. Figure 3F shows that increasing the PBS concentration from 0.5× to 2× significantly reduced the zeta potential of OL from approximately 27 mV to 15 mV. The zeta potential could not be measured at 5× PBS due to the electrical conductivity exceeding the instrument’s limit. This reduction confirms that higher ionic strength compresses the electrical double layer, reduces electrostatic repulsion, and decreases colloidal stability—corroborating the increased coalescence observed under high-salt conditions. These findings indicate that in resisting salt-induced coalescence, OLs play the dominant stabilizing role. Contrary to expectations, the addition of PC did not enhance coalescence stability but instead led to more coalescence events. The mechanism by which PC alters the interfacial structure of OLs will be discussed later. We now turn to the effects of OL and PC on the physical stability of the formed LDs.

### 0.3 The effect of OL/PC-coating on the thin film stability between droplets

After establishing how OLs and PCs influence droplet stability under varying pH and ionic strength conditions, we next investigated their role in the physical stability of LDs. Physical destabilization of emulsions generally occurs through two primary mechanisms: Ostwald ripening and coalescence^27^. In this study, we focus on coalescence, as the generated droplets are highly monodisperse, minimizing Laplace pressure differences that typically drive Ostwald ripening. Moreover, the low solubility of sunflower oil in water further suppresses this process. Consistent with our microfluidic experiments, no evidence for Ostwald ripening was observed.

When two droplets approach each other, a thin liquid film forms between them. The thin film drains due to the Laplace pressure within the droplets and ruptures once its thickness reaches a critical threshold, initiating coalescence. The thin film balance provides a controlled method to investigate the coalescence by mimicking the processes of film formation, drainage, and rupture^28^. Figure 4A illustrates schematically the equipment used in this study. The bike-wheel microfluidic device, where the thin liquid film is generated and controlled, as shown in Supplementary Figure 5, is enclosed within a custom-built pressure chamber connected to a pressure control system and monitored via a camera mounted on an interferometric microscope (see Methods for details). A thick film at mechanical equilibrium is initially formed and then progressively thinned by increasing the oil pressure in the chamber. The applied pressure at the onset of thin film formation is defined as Δ*P* = 0 Pa, while the pressure at rupture is denoted as Δ*P* = *P*_critical_. As shown in Figure 4B, the critical rupture pressure *P*_critical_ is significantly lower for films stabilized by both OLs and PCs (*≈* 58 Pa) compared to those stabilized by OLs alone (*≈* 110 Pa), indicating a weaker interface in the presence of PCs. Microinterferometry images comparing OL-only and OL–PC films at various applied pressures (Δ*P* ) are shown in Figure 4C and D. Due to the small refractive index difference between water and sunflower oil, the images lack the high contrast typically seen at liquid–air interfaces. Nevertheless, both film types exhibit thickness heterogeneity, as evidenced by varying grayscale intensities, contrasting with the uniform appearance of films stabilized by conventional surfactants^29^. Notably, OL-only films displayed heterogeneity at smaller length scales than OL–PC films, particularly at higher Δ*P* , as seen in Figure 4C(iii) and D(iii).

**Figure 4:**
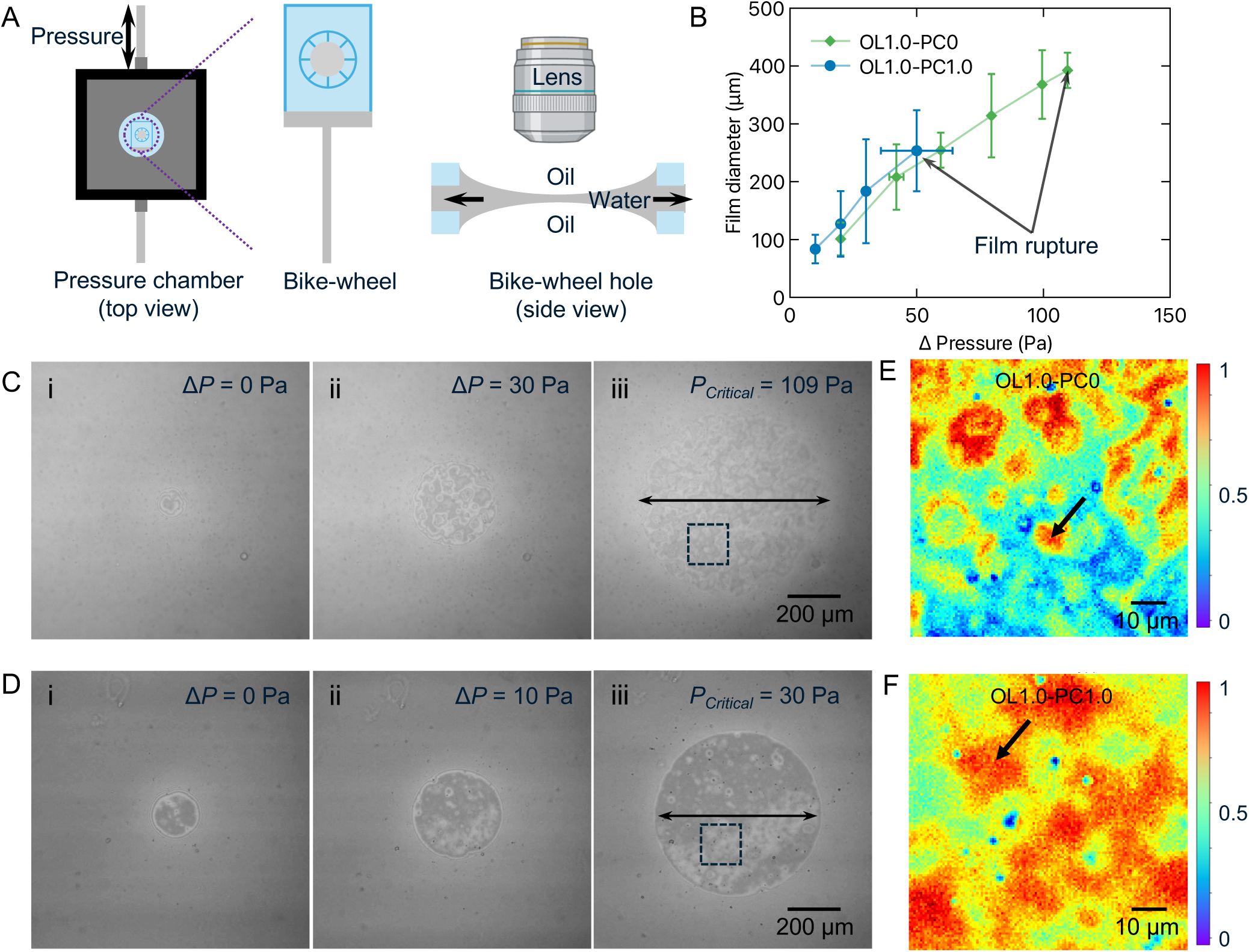
Mimicking coalescence using a pressure controlled dynamic thin film balance. **A** Sketch of the pressure chamber of a dynamic thin film balance and a zoom in of the of the thin water film formed between oil inside the bike-wheel’s hole. **B** Diameter of the film as a function of the applied pressure Δ*P* for films stabilized with OL and with both OL and PC. The last point indicates where the film ruptures. Data are presented as mean ± standard error; *N* = 2 independent experiments in each case. **C** Microinterferometry images of the water film separating two oil phases, with 1 mg/mL OLs dispersed in water phase. i: thin liquid film just formed (Δ*P* = 0 Pa); ii: a thinning film (Δ*P* = 30 Pa); iii: last image recorded before the film ruptured (Δ*P* = 109 Pa). **D** Microinterferometry images of the water film separating two oil phases, with 1 mg/mL OLs dispersed in water phase, and 0.5 mg/mL PCs dispersed in oil phase. From i to iii: thin liquid film just formed (Δ*P* = 0 Pa), a thinning film (Δ*P* = 10 Pa), and the last image recorded just before film ruptures (Δ*P* = 30 Pa). **E** Normalized intensity plots of the area indicated in C-iii, showing high heterogeneity due to the OL aggregates at the interface. Black arrow indicates a typical OL domain at the interface. **F** Normalized intensity plots of the area indicated in D-iii, showing larger domain sizes for the films stabilized with both OL and PC. Black arrow indicates a typical OL domain at the interface.

Sheludko’s equation^30^ is commonly used to calculate the film thickness based on interferometry images of the thin film; however, it has limitations when applied to water films between oil phases stabilized by proteins due to local heterogeneities, the small contrast between the refractive indices of water and sunflower oil, and the difficulty to pinpoint the exact interference order. Instead, the normalized intensity, defined as 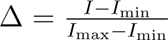, was employed to characterize interfacial heterogeneity. The results suggest that micron-sized OL aggregates form a network at the interface, as shown in Figure 4E, a structure also reported in casein- or cruciferin-stabilized air–water films^22,31^. Upon the addition of PC to the oil phase, the OL domains forming the interfacial network appear larger compared to those in OL-only films, as indicated by the black arrow in Figure 4F. These findings suggest that the lateral phase separation of PC and OL into distinct domains could lead to a loss of percolation in the OL network, thereby contributing to the observed weakening of the interface.

### 0.4 Role of OLs and PCs in modulating droplet interfacial rheology

Our aforementioned experimental observations suggested that the structure of adsorbed proteins at the droplet interface plays a critical role in maintaining the stability of reconstituted droplets. We thus employed interfacial rheology to examine the effects of OL and PC on the surface properties, with particular focus on how PC modulates the network-like structure formed by OLs at the interface and its influence on droplet stability.

We examined the interfacial rheological properties under two conditions: interfaces stabilized solely by OLs and those stabilized by a combination of OLs and PCs. The PC concentration used in the coalescence stability and thin-film rupture experiments could not be applied here, as it caused the droplet to detach from the needle (see Supplementary Figure 6). Instead, a PC concentration of 0.0625 mg/mL was used—the highest concentration that still allowed reliable dilatational measurements. As shown in Figure 5A, the interfacial tension progressively decreases over time in both cases, indicating dynamic adsorption of OLs and PCs at the oil–water interface. Initially, both formulations exhibit a rapid drop in interfacial tension, corresponding to fast adsorption (see also Figure 1C). Over time, the rate of reduction slows and the interfacial tension approaches a plateau, suggesting that interfacial equilibrium is reached. Notably, the interface stabilized solely by OLs consistently showed higher interfacial tension than that stabilized by both OLs and PCs throughout the measurement. Dilatational rheological measurements were performed once the interfacial tension reached a plateau. The resulting interfacial dilatational moduli are presented in Figure 5B. For both systems, the storage modulus (*E^′^* ) was higher than the loss modulus (*E^′′^*), indicating the formation of solid-like interfaces. For the OL-only interface, the storage modulus decreases markedly with increasing strain, reflecting the disruption of the interfacial network at higher amplitudes. In contrast, the OL–PC interface exhibits a weaker strain dependence, with the modulus remaining relatively stable at low strains before gradually decreasing, indicating a more stretchable oil–water interface. Beyond this point, the droplet detaches from the needle, preventing further measurement.

**Figure 5:**
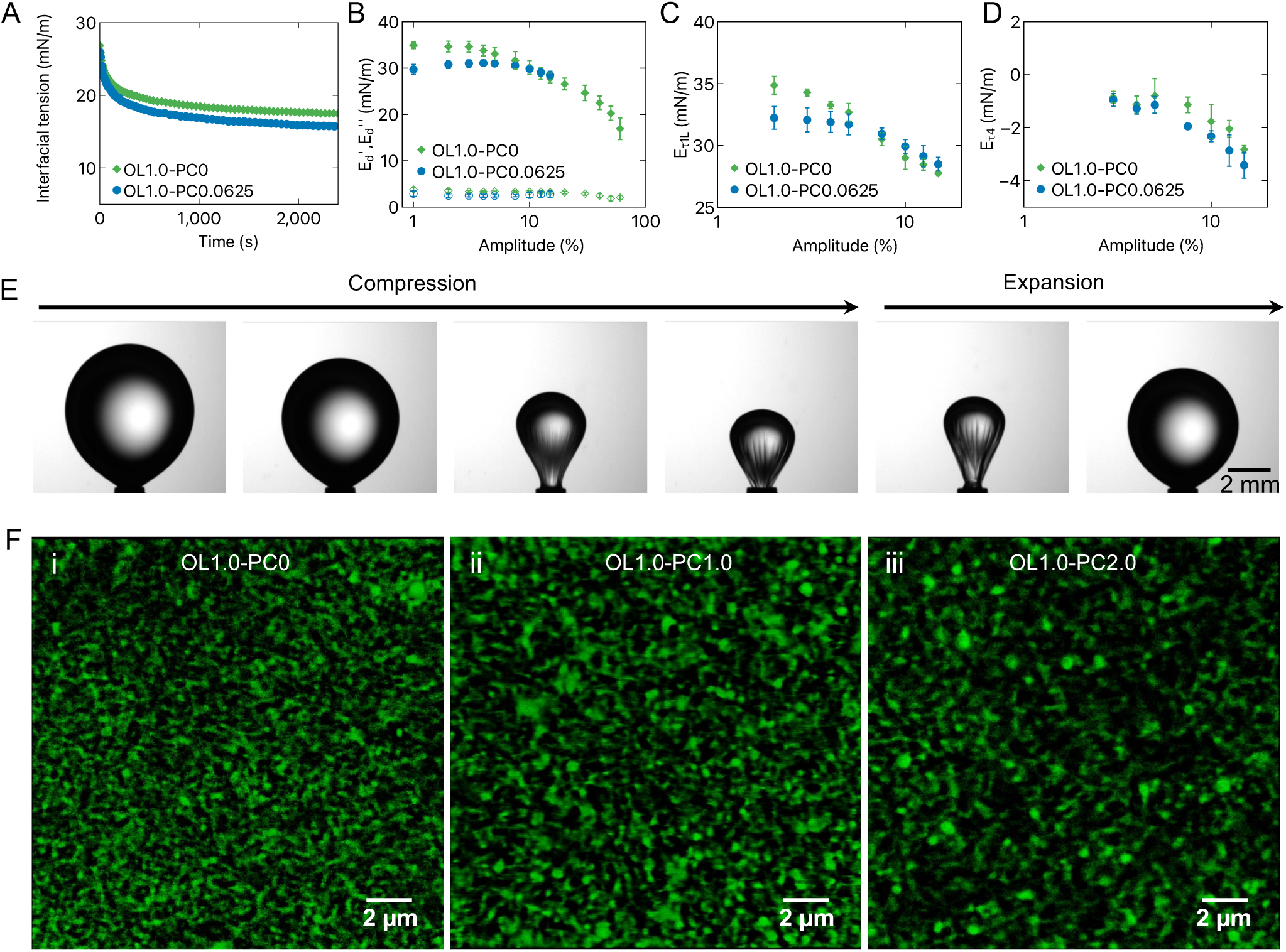
Effects of OLs and PCs on the interfacial structure and rheological properties. **A** Interfacial tension as a function of time measured using a drop tensiometer. Interfaces stabilized by both OLs and PCs exhibited lower interfacial tension compared to those stabilized by OLs alone. **B** Interfacial dilatational modulus as a function of the deformation amplitude. *E^′^* and *E^′′^* represent the elastic and viscous components of the modulus, respectively. Droplets stabilized by OL and PC detach from the needle at amplitudes above 30%, but overall, both *E^′^* and *E^′′^* are higher for OL-stabilized interfaces. **C** Modulus *E_τ_*_1_*_L_*, obtained from odd harmonics, as a function of deformation amplitude. Interfaces stabilized by OLs exhibit higher *E_τ_*_1_*_L_* values at low deformation, while OL–PC interfaces show nearly similar values at larger amplitudes, indicating enhanced interaction between OL and PC under strong deformation. **D** Modulus *E_τ_*_4_ , obtained from the even harmonics, as a function of deformation amplitude, showing that OL-stabilized interfaces consistently exhibit less negative values. **E** Formation of an interfacial layer during large-amplitude oscillations, as evident by reversible wrinkling during compression and and disappearance during expansion, when droplets is stabilized solely by OLs. **F** *i)* STED micrograph showing the interfacial arrangement of OLs in the absence of PCs, indicating the formation of a continuous interfacial network. *ii)* With the addition of PCs, OLs exhibit a similar network structure but with larger domains. *iii)* As the PC concentration increases, OLs become more enriched in larger domains, and the interface appears dominated by PCs.

The results presented (*E^′^* and *E^′′^*) in Figure 5B were calculated based on the first harmonic of the stress responses. For large amplitudes (starting from 5% onward), the stress responses involved signals from higher harmonics, which can be described by a recently proposed general stress decomposition (GSD) method^32–34^ which decomposes the non-linear response into odd harmonics and even harmonics (see Methods for details). The elastic component of *τ*_1_ decomposed from the odd harmonics (i.e., the secant modulus *E_τ_*_1_*_L_* ) can be used to describe the interfacial stiffness. The elastic contribution of *τ*_4_ (i.e., the secant modulus *E_τ_*_4_) obtained from the even harmonics can be used as a measure for the resistance of the interface to changes in surface density. The calculated *E_τ_*_1_*_L_* and *E_τ_*_4_ are shown in Figure 5C and 5D, respectively. The *E_τ_*_1_*_L_* of all interfaces reduced with increased amplitude, indicating a decrease in network stiffness due to the breakdown of interfacial microstructure with increased amplitude. *E_τ_*_1_*_L_* for the interface stabilized only by OLs was higher at smaller amplitudes; however, at larger amplitudes, the interface stabilized by OLs and PCs showed a similar value, indicating that the interfacial response at large deformations is dominated by the OL network. In contrast, the elastic modulus *E_τ_*_4_ (Figure 5D) indicates that surface density differences become apparent at small amplitudes, which is reasonable since the response is governed by the behavior within lipid-rich regions. Figure 5E shows a rising droplet connected to a syringe, subjected to a 50% amplitude compression–expansion cycle. During the initial 50 seconds, liquid is withdrawn from the droplet, resulting in prominent wrinkling of the interface as the droplet shrinks. Upon re-injection, the interface recovered its original shape. The reversible appearance of wrinkles during the deformation cycle suggests the formation of a (visco)elastic skin at the droplet interface^35^. For droplets containing PC, detachment from the syringe needle occurred at a 30% amplitude. At or below this amplitude, neither interface exhibited visible wrinkling during compression, as shown in Supplementary Figure 7. Although the moduli upon addition of PCs are marginally different between 10% to 30%, the differences are striking as the amplitude further increases: droplets stabilized solely by OLs remain attached to the needle, whereas OL–PC–stabilized droplets detach, indicating a much stronger interface when stabilized exclusively by OLs. It should also be noted that the appearance of wrinkles affects the measured dilatational moduli, which are derived from the Laplace–Young equation; therefore, the reported moduli values are apparent^36^. To gain deeper insight into the interfacial network structure, we conducted super-resolution imaging of the collected droplets using STED microscopy. Due to the pancake-like morphology of large droplets in the collection chamber, flat interfacial regions were accessible for imaging. A comparison between confocal and STED micrographs is shown in Supplementary Figure 8. As shown in Figure 5F and Supplementary Figure 9, STED imaging revealed that, in the absence of PCs, OLs formed a continuous interfacial network at the oil–water interface. Upon the addition of PCs, OLs retained this network-like organization but with larger domains, likely resulting from lateral phase separation between PC and OL due to the disruption of lateral interactions between OL C- and N-terminal by acting as a steric spacer, thereby reducing the network density and interface elasticity. Further increase in PC concentration led to OL enrichment within these larger domains, while the remaining interfacial regions appeared to be dominated by PCs. These STED results clearly demonstrate that interfaces stabilized exclusively by OLs possess a denser and a more uniform network, consistent with previous observations on coalescence stability, film rupture behavior, and interfacial rheology.

## DISCUSSION

In this work, we demonstrated that when oleosins (OLs) and phosphatidylcholines (PCs) are used as interfacial stabilizers at the oil–water interface, OLs act as the primary stabilizers of LDs, whereas PCs play a destabilizing role by disrupting interactions among OL C- and N-terminal and thereby weakening the interfacial OL network. We used a microfluidic approach, enabling us to produce and collect monodisperse droplets and further analyze their stability in response to changes in the environment (pH and ionic strength) in a controlled manner. Super-resolution microscopy showed that the adsorbed OLs form an interfacial network and the addition of PCs destabilizes this assembly, resulting in a lower heterogeneity. Indeed, thin film balance experiments showed that the addition of PCs led to a weaker interface, with films stabilized by OLs and PCs rupturing at lower applied pressure than when stabilized by OLs alone. Lastly, interfacial rheology experiments showed that the addition of PCs reduces the elasticity of the interface. Collectively, these results provide clear evidence that OLs, when used as the sole emulsifier, confer enhanced droplet stability, while PCs oppose that effect.

In plant seeds, coalescence-driven changes in oleosome size can disrupt storage organelles, altering lipid and protein accumulation, ultimately affecting germination^37^. Various studies have established a correlation between LD size and the composition of phospholipids and OLs, where larger LDs have been observed under conditions of higher phospholipid levels^38^ or lower OL levels^39,40^. Interestingly, a similar correlation also applies for LDs in plant leaves, hinting that this is not specific to seeds^41^. Our findings demonstrate that the stability of LDs and thus their size can be regulated by modifying the interfacial concentrations of OLs and PCs. Although the results from this work suggest that, in plant seeds, oleosins OLs may function as stabilizers whereas PCs may act as destabilizers, the addition of PC also significantly reduces interfacial elasticity. This reduction enhances interfacial deformability, which is a crucial property in confined environments such as cells. While our reconstituted LDs range from 60 to 100 *µ*m in diameter, most natural LDs are much smaller, from 0.5 to 2.0 *µ*m. Even in specific cases, such as LDs in the mesocarp of oleaginous fruits, where diameters can reach up to 20 *µ*m^38^. The reconstituted LDs in this study are therefore still considerably larger and future work using LDs of sizes comparable to those in nature could help determine whether droplet size also influences stability.

The increasing emphasis on healthy lifestyle and environmental sustainability is driving the demand for natural and sustainable ingredients^42,43^, and this trend is also reflected in the growing adoption of plant-based diets. Emulsifiers are widely used in the food industry, and replacing animal-based or synthetic ingredients with plant-based alternatives such as OL and PC has become of particular interest. The framework developed in this work not only prompts us to reconsider a long-standing assumption that LDs in plant seeds are stabilized cooperatively by interfacial proteins and phospholipids, but also enables the systematic evaluation of the emulsifying properties of proteins or protein–phospholipid mixtures as natural emulsifiers for food applications. In conclusion, we anticipate that this study will not only inspire further research on the effective utilization of natural emulsifiers such as OLs and PCs, but also provide research strategies that open new avenues for investigating how interfacial proteins and phospholipids regulate LDs within plant cells, thereby influencing metabolism and plant development.

## METHODS

### 0.5 Materials

Aqueous solutions were prepared using Milli-Q water with a resistivity of 18 MΩ·cm (Milli-Q, Merck Millipore). Sunflower oil (47123) was purchased from Sigma-Aldrich. Rapeseeds (*Brassica napus*, variety Alizze) were obtained from a European seed producer. Soy PC and fluorescent phospholipid 18:1 Liss Rhod PE were purchased from Avanti Polar Lipids Inc., and were stored at *−*20 *^◦^*C. The thiol-reactive ATTO dye (maleimide, ATTO 643) was obtained from ATTO-TEC GmbH. Microfluidic accessories, including tubing and PDMS couplers, were purchased from Darwin Microfluidics.

### 0.6 Sample preparation

Sunflower oil was purified to remove interfacially active impurities by mixing with Florisil (100–200 mesh, Sigma-Aldrich) at a 2:1 (v/v) ratio. The mixture was covered with aluminum foil and agitated overnight at room temperature using a rotary laboratory shaker. Florisil was then removed by centrifugation, the process repeated two times for 20 minutes each. The purified oil was stored at 4 *^◦^*C until further use. PC dispersions were prepared by dissolving the waxy appearance PC in the purified sunflower oil in an amber vial. The mixture was stirred with a magnetic stirrer for 8 hours until the PC was fully dissolved^44^. OLs were extracted from rapeseed according to previously published protocol^24^. OL dispersions were prepared by dispersing the extracted proteins at a concentration of 1.1 mg/mL in phosphate buffer (137 mM NaCl, 2.7 mM KCl, 8 mM Na_2_HPO_4_, and 2 mM KH_2_PO_4_), pH 2. Tip sonication (Bandelin SONOPULS GM 70) was applied to the solution at 70% power with 50 ms pulses for 5 minutes, and an ice bath was used to prevent temperature increase during sonication. The resulting dispersion was filtered through a 0.45 *µ*m syringe filter to obtain a visually clear, nearly transparent solution. The final protein concentration, approximately 1 mg/mL, was confirmed using the BCA assay as described by Plankensteiner et al.^24^.

### 0.7 Fluorescent tagging of oleosins

Fluorescently labelled oleosins were prepared by using thiol-reactive ATTO 643 dye. First, 6 mM ATTO-643 stock solution was prepared in anhydrous, amine-free DMF and stored in dark conditions. OL dispersion (1 mg/mL) was prepared in PBS at pH 2.0. OL dispersion and ATTO-643 solution were mixed in 1:10 volume ratio and the mixture was incubated at room temperature for 2 hours, protected from light. The incubated sample was filtered through a 0.22 *µ*m syringe filter. Further purification of the resulting conjugate and the removal of unbound dye was performed using size exclusion chromatography (SEC) with a Superdex 200 Increase 10/300GL column (GE Healthcare) on a 1260 Infinity II HPLC (Agilent). Fractions collected were stored at 4 *^◦^*C until further use.

### 0.8 Microfluidic device fabrication

Microfluidic devices were fabricated using a mask-less soft lithography technique. First, SU-8 photoresist (Kayaku) was spin-coated onto a 76 mm diameter silicon wafer (Silicon Materials), with the spin speed adjusted to achieve a uniform target thickness. The microfluidic channel pattern, designed in AutoCAD 2023 (Autodesk), was directly exposed onto the photoresist using a MicroWriter ML 3 photolithography system (Durham Magneto Optics). UV exposure cured the desired pattern, and the uncured photoresist was removed using propylene glycol monomethyl ether acetate (PGMEA, *≥*99.5%, Sigma-Aldrich), leaving cured microstructures on the wafer. To fabricate features of varying heights, the process was repeated twice: SU-8 50 was used to form the inlets, adsorption channels and collection chambers. A mixture of poly(dimethyl siloxane) (PDMS, Sylgard 184, Dow Corning) in a 10:1 base-to-crosslinker volume ratio was poured over the patterned wafer. Air bubbles were removed by vacuum desiccation for 40 minutes, followed by curing at 70 *^◦^*C for 3 hours. The solidified PDMS layer was peeled off, and 0.5 mm diameter holes were punched at the inlets and outlets using a Rapid-Core punch (Darwin Microfluidics). The PDMS channel layer was then bonded to a PDMS-coated cover glass following a 60-second air plasma treatment (PDC 32G, Harrick Plasma). Due to the inherent hydrophobicity of PDMS, which hinders oil/water or water/oil/water emulsion formation, the aqueous-phase-carrying channels must be rendered hydrophilic. Plasma treatment is commonly employed for this purpose^45^. However, its effect is temporary and unsuitable for long-term use. To address this, the channels were filled with Milli-Q water after plasma bonding, which prolonged surface hydrophilicity^17^. Devices treated with 18 W plasma for 60 seconds and stored in Milli-Q water remained hydrophilic and usable for several weeks.

### 0.9 Microfluidic experiments

Oleosins dispersed in 1× PBS, adjusted to pH 2 using HCl, was used as the continuous phase. Sunflower oil mixed with varying PC concentrations served as the dispersed phase. The flow rates of both fluids were independently controlled using a piezoelectric pressure controller (OB1 MK4, Elveflow) with a precision of 12 Pa.

For each coalescence experiment, regions where the droplets were densely packed in a regular hexagonal arrangement were selected to ensure uniform droplet environments. This arrangement was chosen to maintain consistent numbers of neighboring droplets and to assure uniform droplet size and interfacial structure, thereby equalizing the theoretical probability of coalescence for each droplet. The coalescence stability was qualitatively assessed using the coalescence percentage, defined as *N*_coal_ = (*N_t_/N*_0_) × 100, where *N_t_* is the number of coalesced droplets at time *t*, and *N*_0_ is the initial number of droplets. A value of *N*_coal_ = 0 indicates no coalescence, and it increases with the number of coalescence events.

### 0.10 Imaging and post-processing

Bright-field and epifluorescence images were acquired using an inverted fluorescence microscope (Ti2-Eclipse, Nikon) equipped with an illumination system (pE-300ultra, CoolLED) and a sCMOS camera (Prime BSI Express, Teledyne Photometrics). Epifluorescence microscopy was employed to characterize oleosin adsorption during droplet formation at the microfluidic junction. Bright-field microscopy images were used to count droplets and analyze droplet size distribution and coalescence behavior. STED and confocal images were obtained using a stimulated emission depletion microscope (Stellaris DMI8, Leica). The STED microscopy setup consisted of a Leica DMi8 inverted microscope equipped with a white light laser (485–790 nm) and a Leica HC PL APO CS2 86×/1.20 W water immersion objective lens was used for super-resolution characterization. For STED imaging, a 775 nm pulsed depletion laser was used, and fluorescence was detected using HyD detectors. The visualization of thin film balance experiment was performed using a Nikon Eclipse FN1 upright microscope equipped with a Nikon D-LEDI illumination source, a 10× long-working-distance objective (NA = 0.2) and a Hamamatsu ORCA-Flash4.0 V3 CMOS camera. A monochromatic light source at 622 nm is used for reflective imaging. All images were processed using Fiji (ImageJ), MATLAB R2025a or Python.

### 0.11 Oleosin size and Zeta-potential measurement

The zeta potential and size distribution of oleosins were measured at 20 *^◦^*C using a Zetasizer Nano (Malvern Instruments, Malvern, UK). Results were calculated as the average of three independent measurements.

### 0.12 Interfacial tension and interfacial dilatational rheology

The interfacial adsorption behavior of OLs and PCs at the oil–water interface was monitored using a Automatic Droplet Tensiometer (ADT, Teclis, France) over a duration of 1 to 2400 s. A rising oil droplet with a surface area of 30 mm^2^ was formed at the tip of a G18 needle, and its interfacial tension was continuously measured by capturing the droplet contour using a camera. The interfacial tension was calculated by fitting the droplet shape to the Young–Laplace equation. All measurements were conducted in triplicate at 20 *^◦^*C. Following 2400 s of adsorption, interfacial dilatational rheology was performed on the same droplet. Amplitude sweeps were carried out at strain amplitudes ranging from 1% to 60% at a fixed frequency of 0.02 Hz. Five oscillatory cycles were executed per amplitude, and the middle three cycles were further analyzed using the General Stress Decomposition (GSD) method^32^.

### 0.13 General stress decomposition

To characterize the response of interfaces under dilatational deformation^32^, a sinusoidal strain was considered, *ε* = *ε*_0_ sin(*ωt*), where *ε*_0_ is the strain amplitude, *ω* is the angular frequency, and *t* denotes time. In the nonlinear viscoelastic (NLVE) regime, the stress response can be expressed as a Fourier series, capturing the contributions of both even and odd harmonics. This is written as

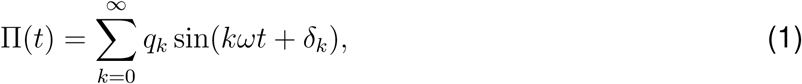

where *q_k_* and *δ_k_*represent the amplitude and phase shift of the *k*^th^ harmonic, respectively. Following the decomposition approach introduced by Yu et al.^46^, the total stress Π(*t*) is separated into four orthogonal contributions based on symmetry transformations:

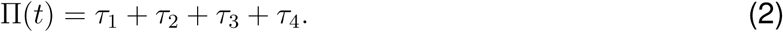

Here, *τ*_1_ and *τ*_2_ represent the odd-harmonic sine and cosine components, respectively, while *τ*_3_ and *τ*_4_ correspond to the even-harmonic sine and cosine terms:

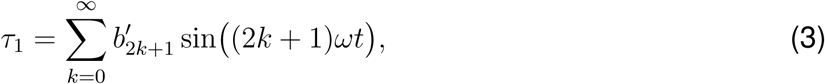

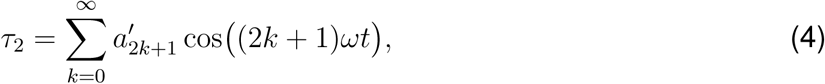

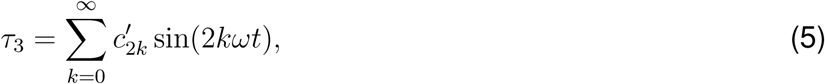

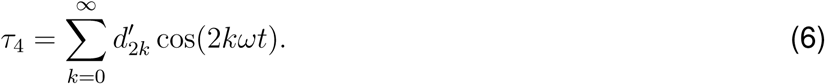

The coefficients 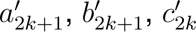, and 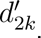 quantify the contributions of the corresponding harmonics. The phase shifts for each harmonic can be obtained from these coefficients. Specifically, for odd harmonics, the phase shift is calculated as 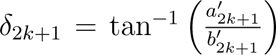, while for even harmonics it is given by 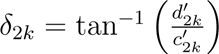. This decomposition enables a systematic analysis of the material response in terms of its linear and nonlinear contributions, capturing the symmetry characteristics of both odd and even harmonic content.

The secant modulus associated with *τ*_1_, denoted as *E_τ_*_1_*_L_*, corresponds to the slope of the line connecting the origin to the stress at maximum extension. It can be obtained from the Fourier coefficients as:

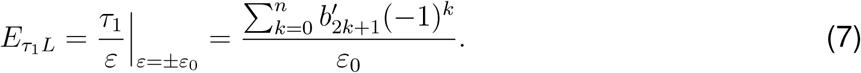

Similarly, for the elastic component *τ*_4_, a secant-type modulus *E_τ_*_4_ is defined as the slope of the dashed line in the *τ*_4_ plot, given by:

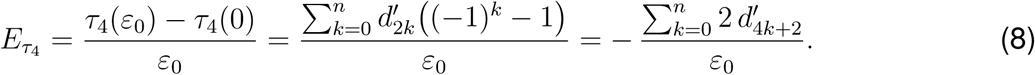

### 0.14 Thin film balance measurements

The bike-wheel chip is a custom-designed microfluidic device fabricated via photolithography on borosilicate glass (Micronit Microfluidics, the Netherlands). The applicability of this method for studying emulsion and foam stability has been demonstrated in previous works^20–22^. The microfluidic device consists of: (i) a diamond-drilled central hole with a radius of 0.5 mm and a thickness of 540 *µ*m, and (ii) 25 radial channels (45 *µ*m wide, 20 *µ*m deep) connected to this central hole, all converging into a larger circular channel, as shown in Supplementary Figure 5. The chip is mounted onto a stainless steel holder using a two-component epoxy adhesive (UHU Plus, Bolton). To enhance contact line pinning between the liquid and glass, the surface is rendered hydrophilic by immersion in a saturated NaOH–ethanol solution, followed by 20 minutes of sonication. Chamber pressure is controlled using a piezoelectric system (OB1 MK3+, Elveflow), with a pressure resolution of 1 Pa. The system is connected to the chamber via rigid PTFE tubing with an inner diameter of 0.1 mm.

## RESOURCE AVAILABILITY

### Lead contact

Requests for further information and resources should be directed to and will be fulfilled by the lead contact, Siddharth Deshpande.

### Materials availability

This study did not generate new materials.

### Data and code availability

Any additional information required to reanalyze the data reported in this paper is available from the lead contact upon request.

## ACKNOWLEDGMENTS

J.v.d.G. acknowledge financial support from the European Union’s Horizon 2020 research and innovation program under the Marie Skłodowska-Curie Actions grant agreement No. 956248. We thank dr. Arjen Bader for the help on the STED microscopy, dr. Lorenz Plankensteiner for preparing the oleosins and the preliminary experimental tests of Jeroen de Heer Kloots, Senna Flameling, and Niyousha Davari.

## AUTHOR CONTRIBUTIONS

X.S., C.V.N., J.v.d.G., and S.D. conceived the research. X.S. and I.A.S. performed the microfluidic experiments. X.S. and X.M. performed the interfacial rheology experiments. X.S., K.A.G. and E.C. performed the thin-film balance experiments. Z.H. performed the SDS-PAGE analysis and fluorescent labeling of oleosins. X.S. wrote the original draft. X.S., J.v.d.G., and S.D. undertook data interpretation, wrote, reviewed, and edited further. All authors have read and agreed to the final version of the manuscript.

## SUPPLEMENTAL INFORMATION INDEX

Figures S1-S9 and their legends

