## Supplementary material for "On the stabilization of plant lipid droplets: Dynamic interplay between oleosins and phospholipids"

[1]Laboratory of Physical Chemistry and Soft Matter, Wageningen University and Research, Stippeneng 4, 6708 WE, Wageningen, The Netherlands
[2]Current affiliation: Laboratory of Food Process Engineering, Wageningen University and Research, Bornse Weilanden 9, 6708 WG, Wageningen, The Netherlands
[3]Laboratory of Physics and Physical Chemistry of Foods, Wageningen University and Research, Bornse Weilanden 9, 6708 WG, Wageningen, The Netherlands
[4]Laboratory of Biobased Chemistry and Technology, Wageningen University and Research, Bornse Weilanden 9, 6708 WG, Wageningen, The Netherlands
[5]Processing and Performance Group, Mechanical Engineering Department, Eindhoven University of Technology, 5600 MB, Eindhoven, The Netherlands
[6]Institute of Complex Molecular Systems, Eindhoven University of Technology, 5600 MB, Eindhoven, The Netherlands

### Suplementary Figures

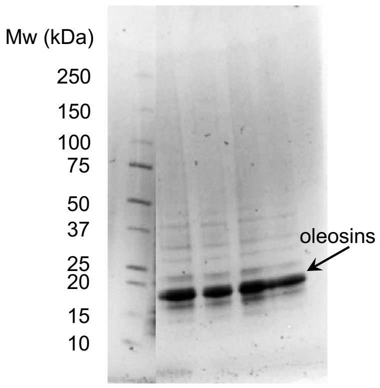


Supplementary Figure 1: **SDS-PAGE profile of oleosins.** A major band at 17 kDa corresponding to oleosins was detected.

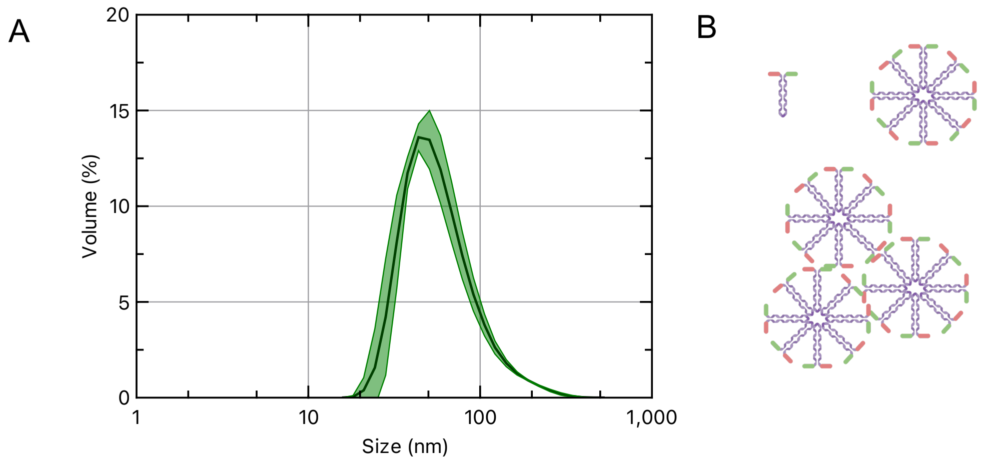


Supplementary Figure 2: **Size distribution of oleosins dispersed in aqueous solution at pH 2. A** Dynamic Light Scattering (DLS) analysis shows a broad size distribution ranging from 20 to 200 nm, with a primary peak at approximately 50 nm. **B** Schematic representation of an individual oleosin, against micelles, and micelle aggregates as suggested from DLS measurements.

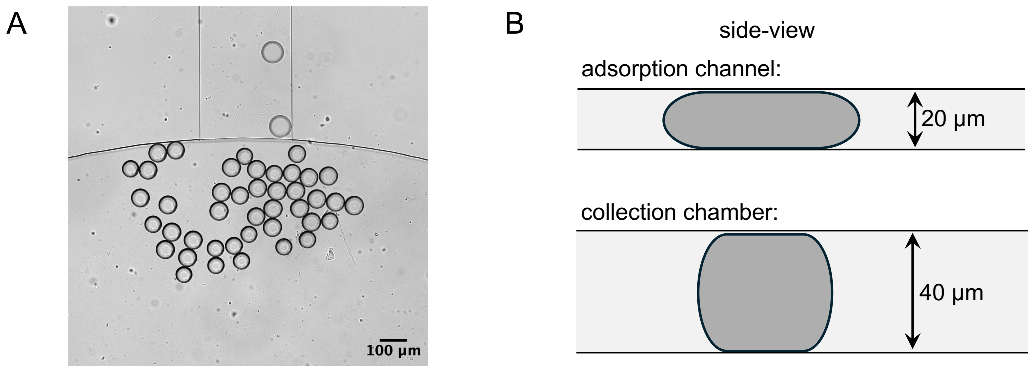


Supplementary Figure 3: **Collection process of lipid droplets.** **A** A multi-height microfluidic device ensures that collected droplets are less squeezed in the collection chamber. **B** Side-view schematic illustrates droplet shape in the channel and the chamber.

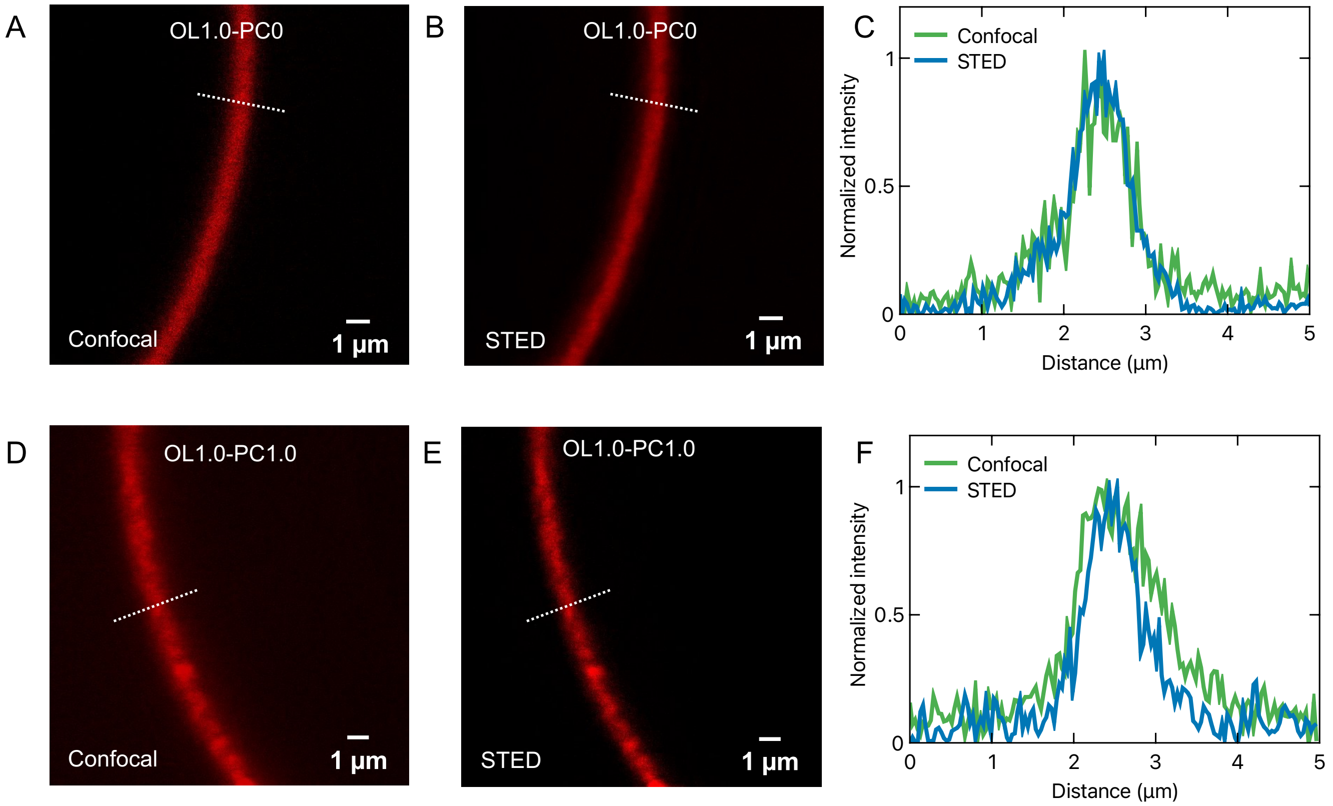


Supplementary Figure 4: **Comparison of confocal and STED micrographs of interfaces stabilized purely by OLs (A–C) and by OLs and PCs (D–F).** Both imaging results demonstrate that STED minimizes in-plane signal interference from background.

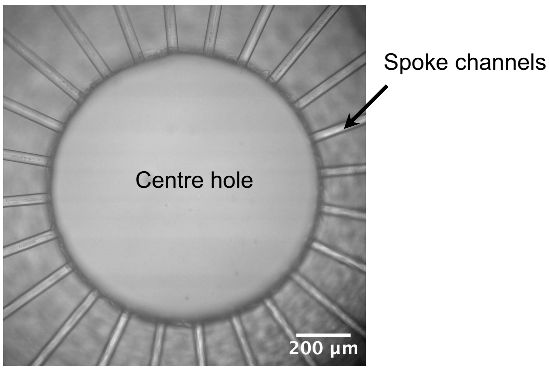


Supplementary Figure 5: **Micrograph of the bike-wheel microfluidic device.** The liquid flows through the spoke channels and forms a film in the central hole. This film is subjected to pressure variations from both above and below.

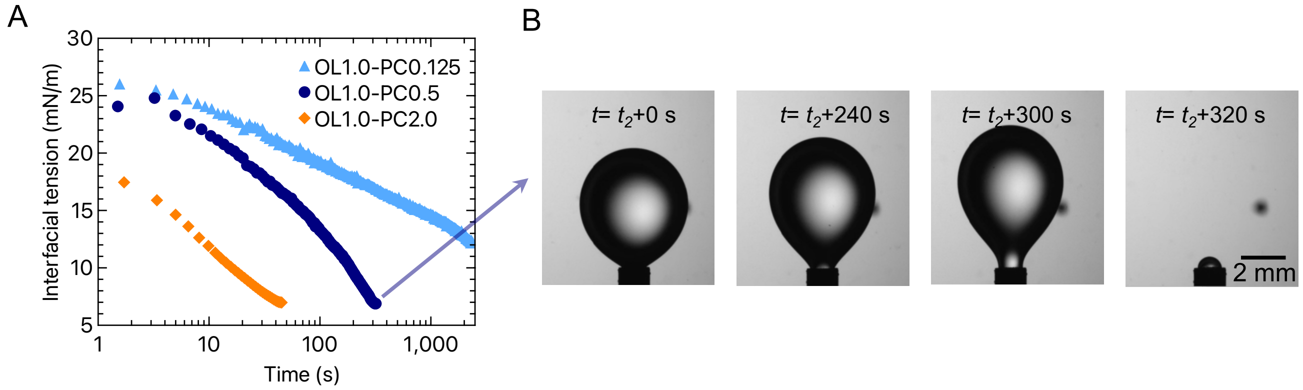


Supplementary Figure 6: **Droplet detachment from the needle at higher PC concentrations.** **A** Interfacial tension as a function of time measured using a drop tensiometer. Droplets stabilized with higher PC concentrations (0.125 mg/mL and above) exhibit reduced interfacial tension and detached earlier from the needle. **B** An example showing the time-lapse of the detachment process.

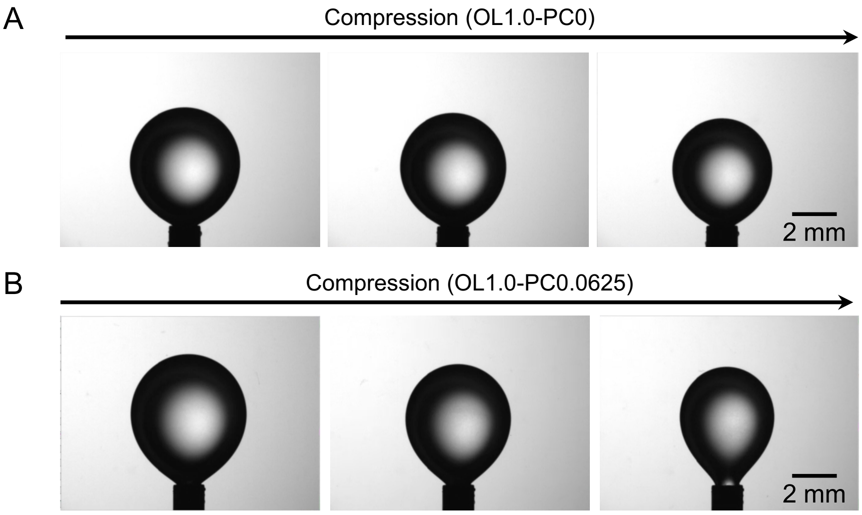


Supplementary Figure 7: **Compression of droplets show no wrinkling in presence of PCs and at small amplitudes.** Droplets stabilized by **A** OL alone as well as **B** OL with PC show no formation of an interfacial layer during oscillation at an amplitude of 30%. In the presence of PC, even at a concentration as low as 0.0625 g/L, droplet detachment occurred when the amplitude exceeded 30%. Consequently, it was not possible to determine whether wrinkling would form at higher amplitudes.

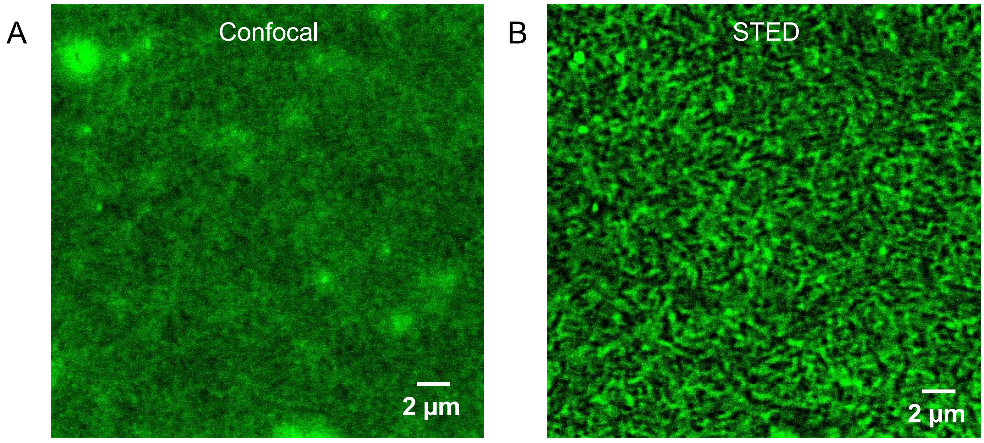


Supplementary Figure 8: **Comparison of confocal (A) and STED (B) micrographs showing the OL distribution at the interface in the absence of PCs.** The STED micrograph clearly shows that the adsorbed OLs form a continuous interfacial network, whereas the confocal images do not reveal the details about the network architecture.

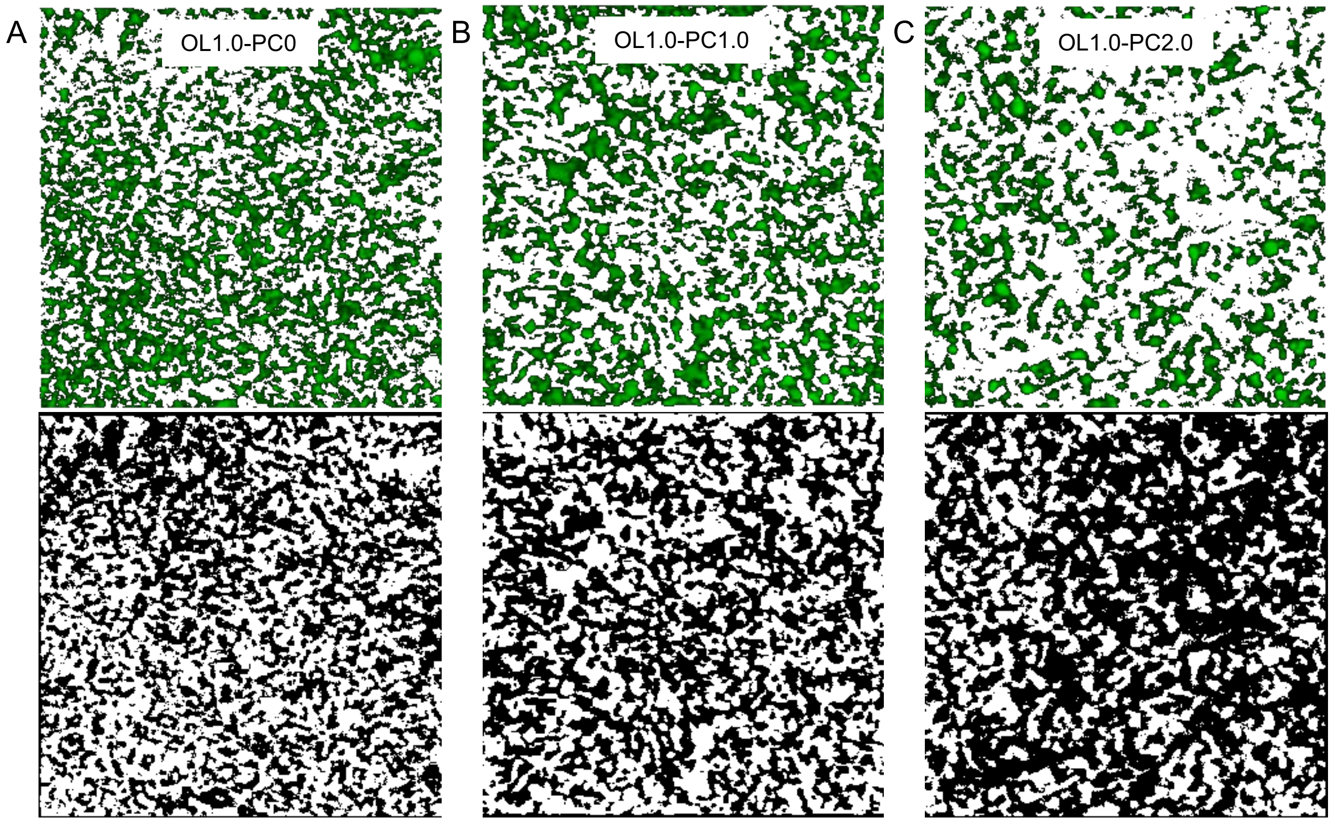


Supplementary Figure 9: **Influence of PC concentration on the structure of interfacial OL network** STED images were processed as single-channel fluorescence micrographs, with the OL signal highlighted in green after threshold-based segmentation (top panel) and non-signal regions displayed in black (bottom panel). The percentage of the green signal showed a consistent decline from $62.24\%$ (**A**) to $57.65\%$ (**B**), ending at $45.45\%$ in **C**.
